# Transcription factor lineages in plant-pathogenic fungi, connecting diversity with fungal virulence

**DOI:** 10.1101/2021.12.16.472873

**Authors:** Evan John, Karam B. Singh, Richard P. Oliver, Kar-Chun Tan

## Abstract

Plant-pathogenic fungi span diverse taxonomic lineages. Their host-infection strategies are often specialised and require the coordinated regulation of molecular virulence factors. Transcription factors (TFs) are fundamental regulators of gene expression, controlling development and virulence in plant pathogenic fungi. Recent research has established regulatory roles for several taxonomically conserved fungal TFs, but the evolution of specific virulence regulators is not well understood. This study sought to explore the representation of TFs across a taxonomically-diverse range of fungi, with a focus on plant pathogens. A significant trend was observed among the obligate, host-associated pathogens, which possess a reduced overall TF repertoire, alluding to a lack of pressure for maintaining diversity. A novel orthology-based analysis is then presented that refined TF classifications, traditionally based on the nature of the DNA-binding domains. Using this analysis, cases of TF over/underrepresentation across fungal pathogen lineages are systematically highlighted. Specific examples are then explored and discussed that included the TF orthologues of Ste12, Pf2 and EBR1, plus phytotoxic secondary-metabolite cluster regulators, which all presented novel and distinct evolutionary insights. Ultimately, as the examples presented demonstrate, this resource can be interrogated to guide functional studies that seek to characterise virulence-specific regulators and shed light on the factors underpinning plant pathogenicity.

## 1. Introduction

Plant-pathogenic fungi have adapted to a wide range of ecological niches. It is now clear that pathogenic lifestyles, broadly defined as biotrophic, necrotrophic or hemibiotrophic, have evolved independently and across distant fungal lineages (Aylward et al., 2017; Möller and Stukenbrock, 2017; Ikeda et al., 2019). As such, the molecular mechanisms underpinning host infections are often specific to a pathogen or its close relatives. In recent years, large-scale comparative genomics and machine learning approaches have demonstrated that the enrichment of particular gene classes can be predictive of a pathogenic lifestyle. This includes carbohydrate-active enzymes, putative effectors, genes related to phytotoxic secondary metabolite (SM) biosynthesis and novel classes that currently lack functional annotations (Collemare and Lebrun, 2011; Pusztahelyi et al., 2016; Plissonneau et al., 2017; Hane et al., 2020; Haridas et al., 2020). For these genes to promote the virulence of a pathogen, their expression requires a coordinated regulation to maximise activity at the appropriate stage of infection (van der Does and Rep, 2017). Hence, the evolution of virulence encoding genes simultaneously requires the pathogen to evolve appropriate regulatory systems.

Transcription factors (TFs) regulate diverse aspects of fungal development and virulence, targeting either a small number of genes or by exerting broad control as ‘master’ regulators. Conserved TFs originally characterised in model saprophytes are now well-studied in plant-pathogenic fungi (Tan and Oliver, 2017; van der Does and Rep, 2017; John et al., 2021). However, there is extensive variation in the number of TFs belonging to different structural families across the fungal taxa (Park et al., 2008; Todd et al., 2014; Shelest, 2017) and the role of most remains undefined. It is therefore anticipated that a functional exploration of the TFs unique to a fungal lineage or lifestyle will reveal novel regulators that distinguish a pathogen from a non-pathogen. Already, TF gene duplications that are directly implicated in fungal virulence have been described in several lineages of *Fusarium oxysporum*, the causal agent of vascular wilt on a wide range of plants (de Vega-Bartol et al., 2010; Niño Sánchez et al., 2016; van der Does et al., 2016). On the other hand, broadly conserved regulators in plant-pathogenic fungi, such as the Velvet TFs, are absent in saprophytic yeasts (Bayram and Braus, 2012; Calvo et al., 2016). Hence, in both their presence or absence, TFs are implicated in the distinct regulatory systems that have evolved to control plant-pathogenic lifestyles.

Random mutagenesis or high-throughput gene knockout studies are traditional approaches that have been used to identify novel virulence-regulating TFs. For example, VdFTF1 in *Verticillium dahliae*, which regulates the expression of secreted virulence factors during infection on cotton, was identified by screening a random mutagenesis library (Zhang et al., 2018). On the other hand, a genome-wide TF knockout screen was undertaken for *Fusarium graminearum*, which identified several virulence regulators otherwise dispensable for saprophytic growth (Son et al., 2011). However, the generation and screening of fungal mutant libraries in this fashion can be labour intensive. An approach that makes use of the annotated fungal genomes now available to disentangle regulators unique to a pathogenic lineage or lifestyle presents a powerful stand-alone approach, or as an initial step to help guide functional TF investigations.

A novel analysis is presented here that simultaneously assesses the TF repertoires (“TFomes”) from 100 phytopathogenic fungi alongside 20 representative saprophytes/symbionts for comparison. The TFs were refined into orthogroups; the smallest definable set of proteins where all their orthologues are included (Emms and Kelly, 2019). In doing so, the relationships between TFs across plant-pathogenic fungi could be systematically assessed. The distinct aims were to A) explore TF conservation, expansion and loss associated with plant-pathogenic lifestyles and B) provide a novel resource for tracing the evolutionary trajectory of TFs correlated with virulence in distinct fungal lineages. Using the orthogroups defined in the analysis, the evolutionary lineages of several established fungal-virulence regulators are explored in detail. These presented examples of TF conservation, expansion/diversification and loss in key pathogen lineages and also provided novel insights upon which future analyses can build.

## 2. Methods

### 2.1. Compiling fungal TFomes

Non-redundant fungal proteomes were sourced for 100 plant pathogens, 10 saprophytes and 10 symbionts from UniProt (release 2020_05) (Bursteinas et al., 2016) or MycoCosm (Nordberg et al., 2014) and independent platforms where the proteomes were available with higher coverage assemblies (**Supplementary item 1**). Pathogens were selected from diverse fungal taxa of significant research interest (Pedro et al., 2016; Urban et al., 2020). The saprophytes and symbionts included both close and distant pathogen relatives across the taxa. Organisms were assigned a five letter organism ID (ORGID) for brevity (**Supplementary item 1**), adapted from their NCBI Taxonomy database naming convention (Schoch et al., 2020). General lifestyles (biotroph, hemibiotroph, necrotroph, saprophyte or symbiont) and plant host-associations/dependency (i.e. obligate, facultative or not associated) were assigned based on descriptions from the published proteome sources and/or literature covering the matter (Möller and Stukenbrock, 2017; Hane et al., 2020; Haridas et al., 2020).

Interpro protein-domain annotations were also downloaded with the respective fungal proteome sources. Interproscan (release 82.0) was used to annotate proteomes from the independent servers where these were not available (Blum et al., 2020). TFs were selected from the annotated proteomes by cross-referencing a list of Interpro domain accessions that represent sequence-specific DNA-binding domains (DBDs) (**Supplementary item 2**). Unique protein identifiers were assigned to each TF following the format ‘ProteinID | ORGID’ where the ‘ProteinID’ was derived from the downloaded proteome source.

### 2.2. TFome regression analysis

The total occurrences of each TF domain across the annotated fungal proteomes (which included cases where multiple domains exist within a single protein) were counted using a custom bash script. Linear regressions were then produced to model the association between proteome size and TFome size for the entire set of 120 fungi, as well as for the lifestyle and host-association sub-groups. The data analysis, formatting and plotting was undertaken in R (version 4) using the ggplot2, ggpubr, rstats, rstatix and emmeans packages (R Core Team, 2020). Pearson’s correlation coefficients (Mukaka, 2012) were calculated for each linear regression and used as the test statistic to assess whether the association was significant (P < 0.05). A covariance analysis (ANCOVA) was also undertaken to test whether the regression-slopes (i.e. the rate of TFome size increase relative to the proteome size) were different between the lifestyle or host-association groups (Bonferroni Padj < 0.05).

### 2.3. TF orthology analysis and species-tree inference

An orthology analysis for the entire TF set compiled from the TFomes was undertaken using OrthoFinder (release 2.4.0) (Emms and Kelly, 2019), with MMseqs2 (Steinegger and Söding, 2017) invoked for the sequence search stage. The TF-distance based species-tree (which included here several sub-species taxonomic rankings), produced as part of the OrthoFinder analysis by STAG (Emms and Kelly, 2018), was manually rooted at the most distantly related pathogen *Synchytrium endobioticum* (SYNEN), inferred from the phylogeny documented in a previous study (Choi and Kim, 2017). OrthoFinder was subsequently invoked using the ‘-s’ option and supplied the rooted species-tree to resolve the orthogroup ‘protein-trees’. The protein-trees were built using the DendroBLAST distance measure (Kelly and Maini, 2013) as part of the OrthoFinder analysis. Tree visualisations and formatting were subsequently undertaken using iTOL (Letunic and Bork, 2019). NCBI taxonomic rankings for each respective phylum, class, order, family and genus were then sourced using Taxize (version 0.9.99) to demonstrate the clades in the TF-distance based species-tree corresponded with fungal taxonomies (Chamberlain et al., 2020).

Fungal TF-orthogroup datasets were ordered as per the leaf-node order derived from the species-tree (beginning with the root at *S*. *endobioticum*) for subsequent visualisation and analyses. Data formatting and the production of heatmaps was undertaken in R (version 4) using the phylogram, complex heatmap, ggplot2, ggpubr and the rstats packages (R Core Team, 2020). A hierarchical clustering analysis was conducted to group the TF orthogroups based on their relative counts across the 120 fungi analysed. The Z-scores were first calculated similarly to a previous TF analysis (Charoensawan et al., 2010a), which represented a clustering measure independent of the differences in orthogroup sizes. Clustering distances were calculated based on Pearson’s correlation and the orthogroups were clustered using the Average linkage function.

### 2.4. Independent analysis of established fungal-virulence regulators

Orthogroups harbouring established virulence regulators were determined by cross-referencing the respective ProteinIDs with experimentally validated TFs from the scientific literature (John et al., 2021). TFs of evolutionary interest were compiled by inspection of the respective orthogroup protein-trees. A multiple-sequence alignment was then undertaken on the selected protein sequences using Clustal Omega (Sievers and Higgins, 2018) and conserved regions corresponding to Interpro-domain annotations (Blum et al., 2020) were assessed across the protein set. These alignments were then used to produce Neighbour-Joining trees with the Jukes-Cantor distance measure in Geneious Prime (version 2021) as a complementary method to assess the orthogroup clade-topologies.

## 3. Results & Discussion

### 3.1. Correlations between TFome and proteome sizes in plant pathogenic fungi

Fungal TFomes were defined as the set of proteins harbouring at least one sequence-specific DBD in line with previous analyses (Park et al., 2008; Shelest, 2008; Todd et al., 2014; Shelest, 2017). The list of corresponding Interpro domains was updated (**Supplementary item 2**) from the most recent analysis (Shelest, 2017) by incorporating 16 additional sequence-specific DBDs such as the Velvet domain (IPR037525), the high-mobility-group box domain (IPR009071), the APSES/Swi6 domain (IPR001606) and the Sant/Myb domain (IPR001005). Conversely, two domains were excluded that represented single-stranded nucleic-acid binding proteins (IPR012340 and IPR001878). Hence, the 120 fungal TFomes were assembled from refined TF-annotation criteria. Their relative size across each of the 120 proteomes was first assessed to gauge any broader trends in the TF-regulatory capacity (**Fig. 1**). These ranged from 189 TFs identified (for *Blumeria graminis* f. sp. *tritici*; BLUGT) to 1,493 (for the *F. oxysporum* biocontrol strain Fo47; FUSO4). The average TFome size was 473 from an average proteome size of 12,355. This corresponded to a median of 437.5 and 12,296.5 respectively. The TF fraction of the proteomes ranged between 1.2% (for *Puccinia triticina*; PUCTR) to 6.9% (for FUSO4) with an average of 3.9%. The highest proportion of TFs was generally observed among the *Fusarium* lineages, while rust pathogens such as *Melampsora* and *Puccinia* spp. shared a relatively low TF content. Overall, a moderate-linear correlation (*R* = 0.58) was observed between TFome size and proteome size. A previous study that assessed a broad range of eukaryotes reported a stronger correlation (*R* LJ=LJ0.78) (Charoensawan et al., 2010b). This suggested TF-content variation was higher among the fungi assessed in this study.

**Fig. 1.**
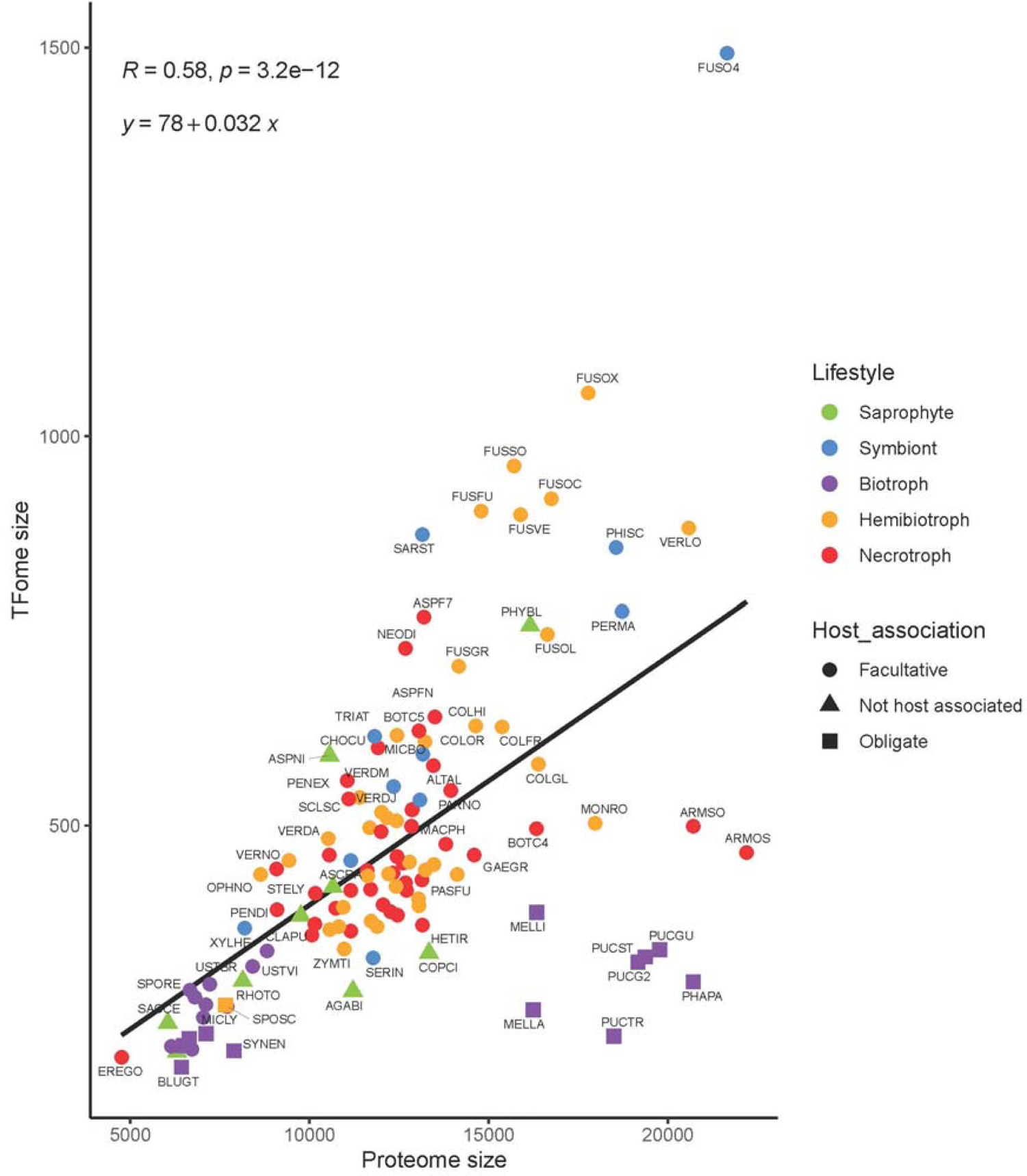
Fungal transcription factor content (TFome) relative to proteome size TFome vs proteome sizes for the 120 fungi used in this analysis. The five-letter fungal ORGIDs are used to label the respective points which provide an indication of the respective lifestyle and host-association (**Supplemental item A**). The correlation coefficient (*R*) and P-value (*p*) used to test for any association (P < 0.05) and linear regression (*y*) for the relationship are presented at the upper-left of the plot.

The TF content of fungal sub-groups, defined by their lifestyle or association with plant hosts, were then compared to explore any distinct trends relevant to phytopathogenicity. Linear correlations were observed for TFome vs proteome counts from each pathogenic (i.e. biotrophic, hemibiotrophic or necrotrophic) and non-pathogenic (i.e. saprohphytic or symbiotic) fungal lifestyle (**Fig. 2**). No significant differences were observed across the linear models derived from these groups. This was not the case when fungi were grouped based on their host-associations (i.e. facultative, obligate or not host-associated). The TF content of obligate-host fungi relative to the proteome size was significantly lower than facultative pathogens and non-host saprophytes (**Fig. 2**). These were largely represented by the rust and mildew pathogens from the Pucciniales (1.7% average TF fraction) and Erysiphales (3.2% average TF fraction) fungal orders. This observation provides evidence that a restricted regulatory capacity exists where fungal pathogens are intimately connected with their host for survival. Conversely, fungi which occupy broader ecological niches are likely to encounter diverse selection pressures that maintain TF diversity. Some outliers to these trends were observed in distinct fungal lineages. For example, the average TF fraction in the Agaricomycetes class, which included several facultative necrotrophs and non-pathogenic saprophytes, was just 2.5%. TF-content comparisons within lineages are therefore necessary to resolve the specific host-lifestyle differences that have evolved.

**Fig. 2.**
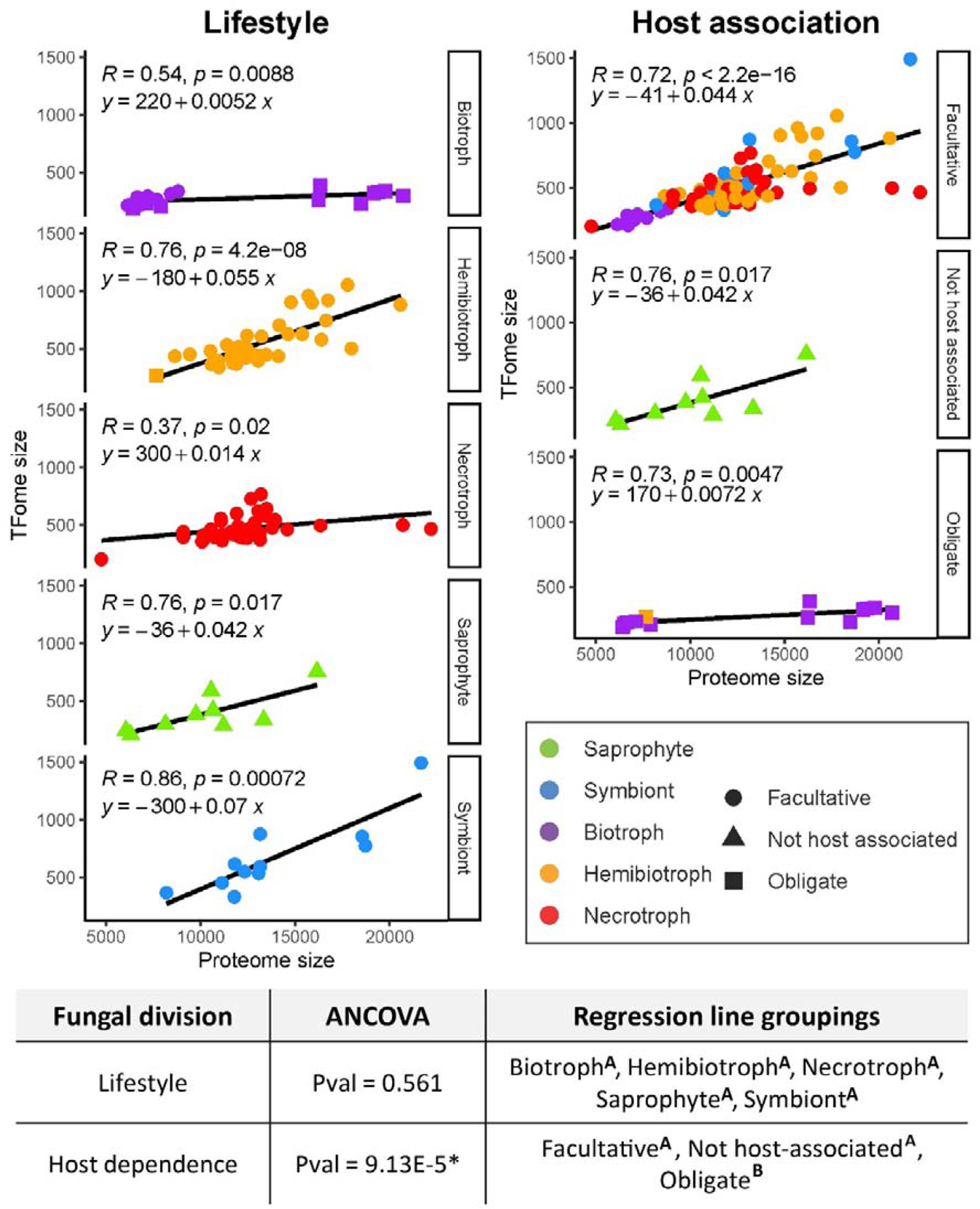
A comparison of the transcription factor proportion of fungal proteomes grouped by lifestyle or host-association TFome vs proteome sizes are depicted along with the correlation coefficients (*R*) and P-values (*p*) for the respective linear regressions (*y*) in each plot. The panel on the left depicts the relationships by lifestyle and the right panel by host-association for the 120 fungal proteomes assessed in this study. The bottom table summarises where a statistically significant difference (Pval < 0.05)* exists across the regression rates determined by ANCOVA. Superscript letters A or B indicate where individual differences occur within the groups.

### 3.2. Lineage-specific TF orthogroup expansions among plant-pathogenic fungi

A cursory look at the total set of fungal TFs classified by their DBD-based families (**Supplementary item 2**) revealed that the Zn2Cys6 and C2H2 zinc finger domains were the most prevalent among the 120 fungal species/strains, which paralleled previous analyses (Todd et al., 2014; Shelest, 2017). A total of 32,023 Zn2Cys6, 11,436 C2H2, 4,777 homeodomain, 3,168 bZIP and 2,932 CCCH-type domains represented the top five among the 70 TF families where at least 10 instances were identified overall. The large size of such families motivated an orthology analysis with the aim to better classify the TFs represented in distinct fungal lineages. This resolved the 56,754 fungal TFs into 855 orthogroups (**Supplementary item 3**), which presented a large refinement of the DBD-based classification system. The TF-distance based species-tree derived from the fungal orthogroups was then compared with the fungal taxonomic clade-architecture to assess the overall accuracy of the analysis. Consistent topologies were observed (**Fig. 3**; **Supplementary items 4 & 5**) which suggested that sequence divergence within the TF orthogroups correlated well with species divergence across fungal lineages.

**Fig. 3.**
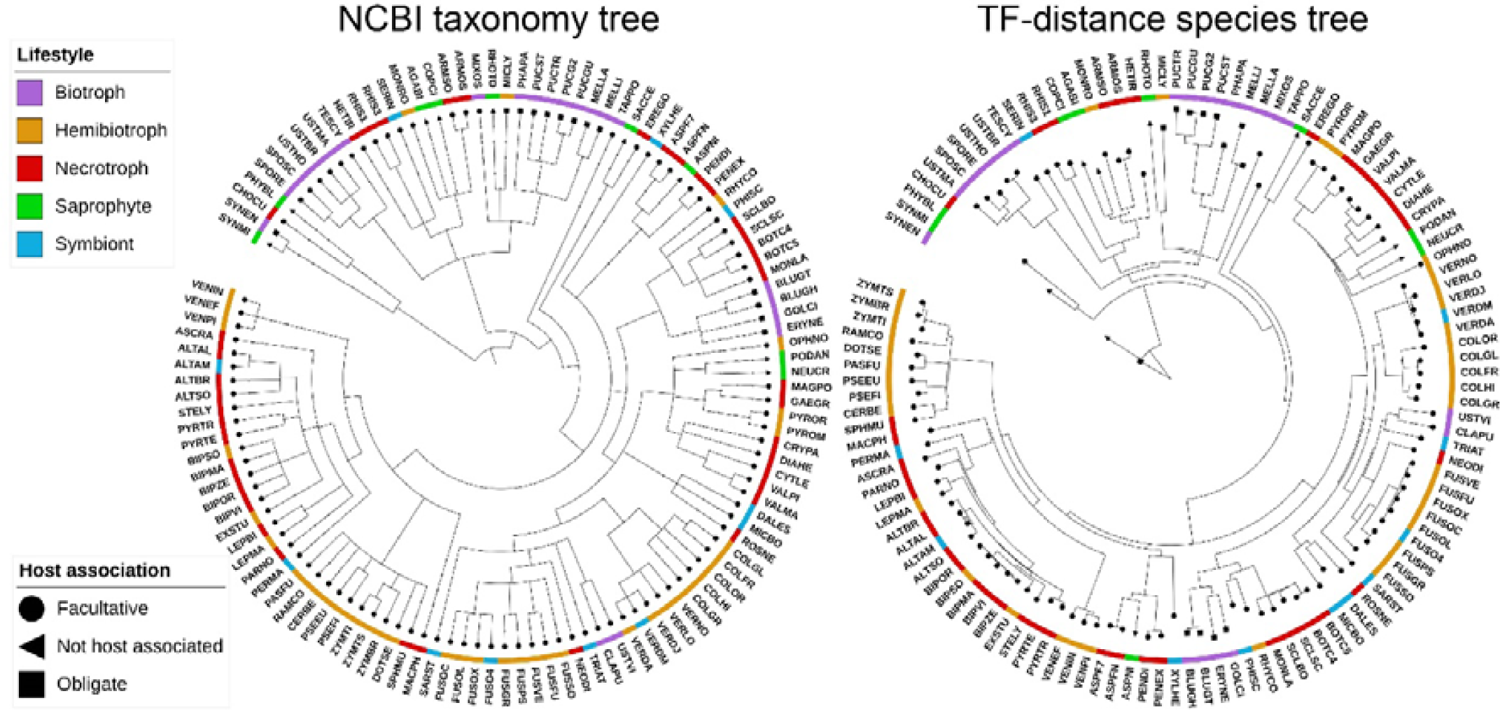
Fungal taxonomy relative to the transcription factor (TF)-distance based species-tree Clade architectures derived from NCBI taxonomic divisions (node dots) are presented in the left tree, demonstrating shared topologies with the TF-distance based species-tree. Fungal ORGIDs are used at branch tips due to space constraints (**Supplementary item 1**). The corresponding trees using the full species names are available for higher resolution viewing as **Supplementary item 4 & 5**.

The TF orthogroups were then clustered based on their normalised counts across the fungal species/strains (**Supplementary item 6**). This sought to assess the extent to which the respective TFs are expanded or specific to distinct pathogen lineages, a potential indicator for virulence-associated selection. Inspection of the corresponding heatmap (**Fig. 4**) highlighted widespread cases of overrepresentation down to the genus, species or sub-species level. While the relevant consequences in the capacity for individual fungi to regulate virulence will need to be functionally assessed, the analysis demonstrated evolutionarily recent shifts are traceable to specific TF lineages.

**Fig. 4.**
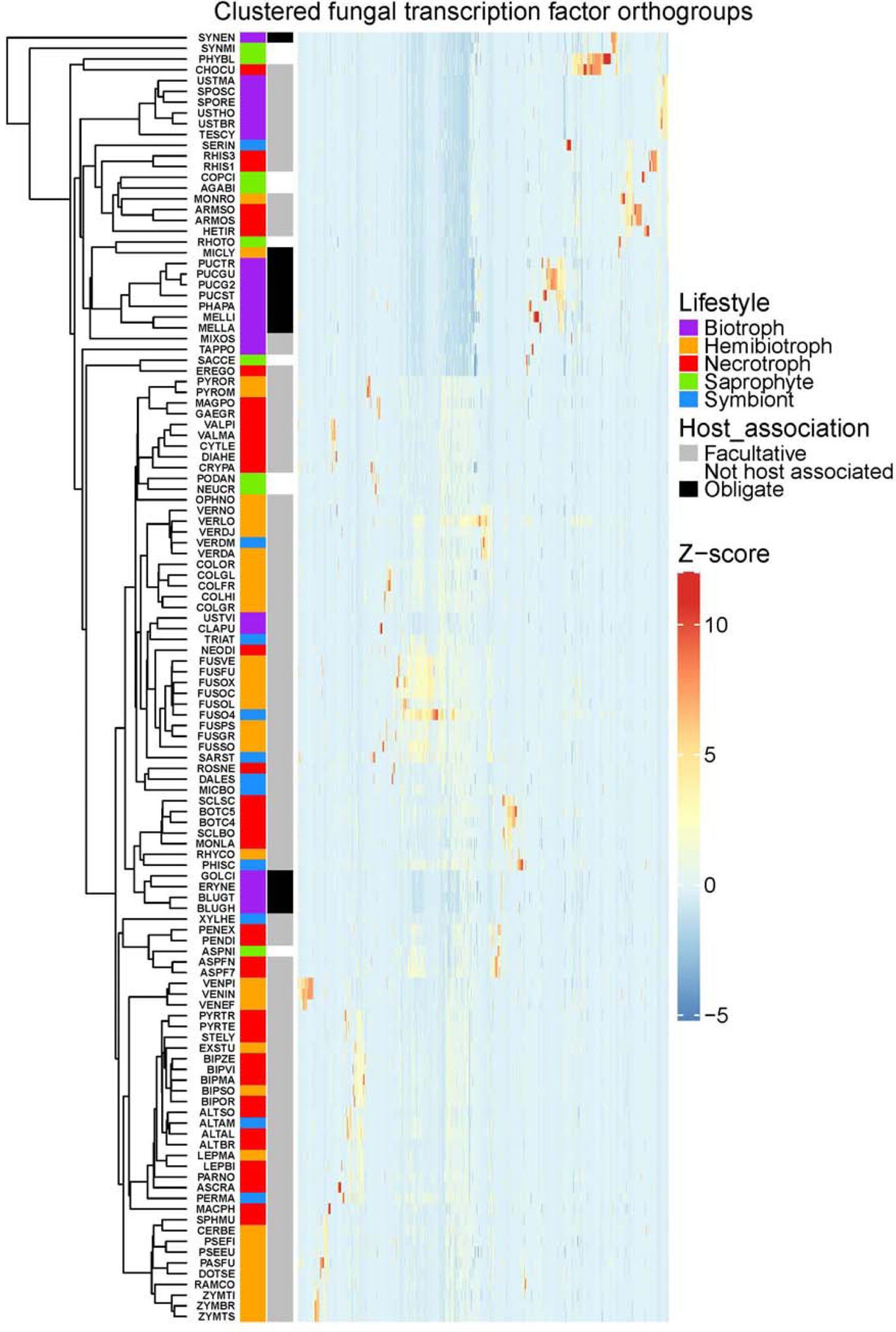
Relative proportion of transcription factors (TFs) within orthogroups across the 120 fungi assessed in this study A heatmap depicting TF expansion based on relative counts (Z scores) for each of the 855 orthogroups defined in this analysis (arrayed on the X axis and clustered using the Euclidean distance measure). Fungal ORGIDs are listed on the Y-axis based on the TF-species-tree order depicted in **Fig. 3**. Coloured markers adjacent to ORGIDs indicate the respective fungal lifestyle and plant host-association. The TF counts used to produce this figure are provided (**Supplementary item 6**) along with a high-resolution version of the heatmap that details the respective IDs for the clustered orthogroups (**Supplementary item 7**).

### 3.3. Regulators of virulence in conserved and recently evolved TFs

The breadth of the 855 orthogroups defined in the analysis constrained the extent to which individual fungal lineages could be assessed in detail. Instead, the evolutionary trajectories for several established virulence regulators are explored here in the context of their respective orthogroups to provide examples where novel insights can be gained into fungal plant pathogenicity. This makes use of the orthogroup ‘protein-trees’ produced from the analysis which allow the predicted origins of TF duplication and/or loss to be traced across fungal lineages. The respective protein-trees for each orthogroup (OG0000000 > OG0000854) are provided as supplementary material (**Supplementary item 8**). The virulence regulators explored here include TFs which regulate either a broad or restricted set of genetic pathways, as well as those that are conserved across fungal taxa and those where orthologues are difficult to trace.

#### 3.3.1. Pf2-orthologues underpin virulence within a lineage of carbohydrate metabolic regulators

Orthogroup OG0000017 comprises 370 annotated TFs that belong to the Zn2Cys6 DBD family. These are represented across the Ascomycota, in most cases with multiple TFs identified for each fungus. This includes MAL13, ZNF1, MAL33 and YGR288W characterised in *Saccharomyces cerevisiae* that target either the aerobic and/or anaerobic pathways regulating carbohydrate metabolism (Akache et al., 2001; Tangsombatvichit et al., 2015). Analysis of the OG0000017 protein-tree revealed these four TFs were adjacent to each other in a single clade, suggesting they are paralogues in the context of this orthogroup (**Supplementary item 9**). Further inspection of the protein-tree revealed another clade harboured the AmyR regulator (ANIA_02016) from the filamentous saprophyte *Aspergillus nidulans*. AmyR is known to play a fundamental role in starch/polysaccharide hydrolysis (Tani et al., 2001; Nakamura et al., 2006). The common link to carbohydrate metabolism for these regulators is also conserved in a separate clade that is of significant interest from the perspective of phytopathogenicity. This encompasses Pf2 (Pleosporales spp.) and several other virulence-regulating TFs that include ART1 (*Fusarium* spp.), MoCod1 (*Magnaporthe oryzae*), PdeR (*Botrytis cinerea*) and Zt107320 (*Zymoseptoria tritici*) (Cho et al., 2013; Chung et al., 2013; Oh et al., 2016; Jones et al., 2019; Habig et al., 2020; Han et al., 2020). The analysis suggested each of these virulence-regulators are traceable as direct taxonomic orthologues to *Neurospora crassa* Col-26 (**Supplementary item 9**). Recent studies have established the important role of Col-26 in polysaccharide metabolism and plant cell-wall degradation (Xiong et al., 2017; Li et al., 2021).

It was intriguing that the Pf2/Col-26 monophyletic clade within OG0000017 did not include TFs from the Eurotiomycetes or Saccharomycetes, such as AmyR or MAL13, but is conserved among the other Ascomycete lineages. A protein sequence alignment/phylogenetic tree was therefore built from the functionally-characterised MAL13, AmyR and Pf2-orthologues to highlight the common and divergent features (**Fig. 5**). A relatively high degree of sequence conservation was observed at the Zn2Cys6 DBD (IPR001138) and the ‘Fungal transcription factor’ domain (IPR007219) that are present across the proteins. However, a region unique to the Pf2-orthologues was identified at a sequence centred on the C-terminus of the Zn2Cys6 DBD (**Fig. 5**), suggesting differentiation relevant to DNA-binding. Previous studies that screened Zn2Cys6 gene-deletion-mutant libraries in both *M. oryzae* and *F. graminearum* (Son et al., 2011; Lu et al., 2014) support a distinct role for the Pf2-orthologues. In these analyses no obvious virulence-defects were associated with any of the 12 other TFs identified in OG0000017, which suggested the Pf2-orthologues (MoCod1 and FgART1 in these species) are the prominent regulators in this orthogroup.

**Fig. 5.**
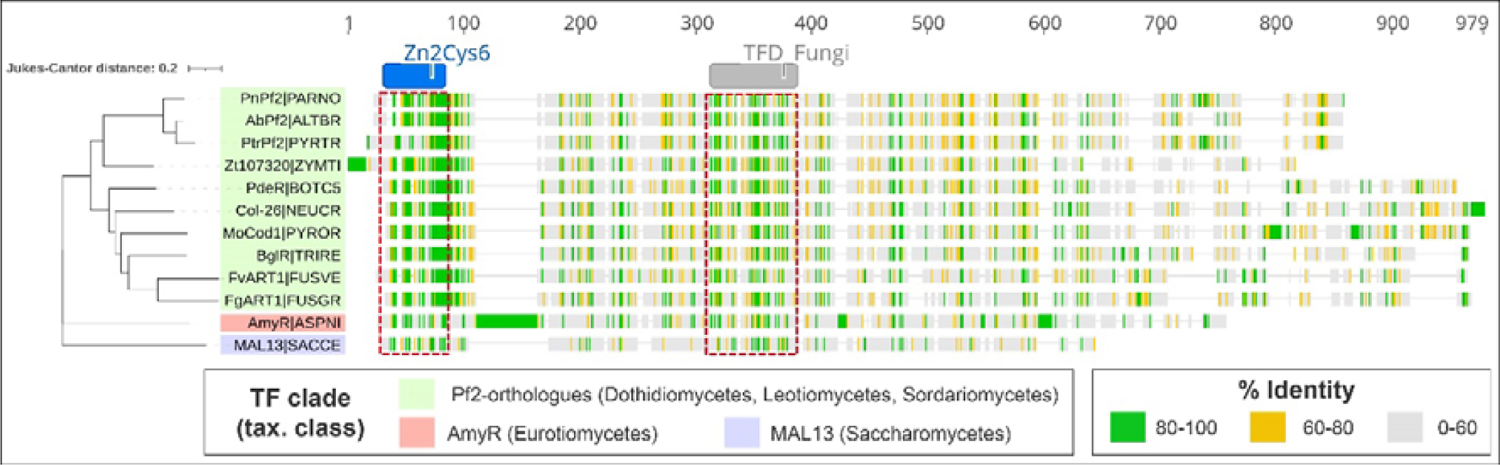
Characterised Pf2-orthologues in relation to established carbohydrate regulators A protein-sequence alignment is presented with a neighbour-joining tree depicting sequence divergence between Pf2-orthologues (highlighted in green) and the *A. nidulans* AmyR and *S. cerevisiae* MAL13 proteins that were identified in orthogroup OG0000017. The protein names correspond to those reported in the relevant literature followed by the five-letter ORGIDs used in this study for the respective fungi. The alignment presents the % of residue identity across the aligned loci with gaps being ignored. The regions corresponding to the Zn2Cys6 (IPR001138) and the ‘Fungal transcription factor’ (IPR007219) Interpro domains that were identified are presented above the alignment. Red boxes span the sequences where the respective domains are detected.

The analysis of the OG0000017 orthogroup provided clarity for the relationship between Pf2-orthologues and their probable evolutionary origin from a lineage of carbohydrate/polysaccharide metabolic regulators. These processes are fundamental for plant pathogens that exhibit a necrotrophic phase of their lifecycle for the acquisition of nutrients from their respective hosts. A question that remains will be to establish the unique features that evolved for Pf2-orthologues which explain the distinct and conserved role reported in fungal virulence.

#### 3.3.2. Evidence of domain architecture variation in Ste12 TFs

Ste12 TFs are downstream targets of mitogen-activated protein kinase (MAPK) signalling that control expansive hyphal growth (as opposed to yeast-like budding growth) and the pheromone response (sexual development) in *S. cerevisiae* (Rispail and Di Pietro, 2010; Zhou et al., 2020). An analogous function has been attributed in fungal phytopathogens, where Ste12 TFs play a significant role in plant virulence that is manifested through the regulation of invasive hyphal growth (Wong Sak Hoi and Dumas, 2010; John et al., 2021). Interestingly, Ste12 orthologues in filamentous fungi also harbour C2H2 DBDs located in the C-terminal region of the protein (Wong Sak Hoi and Dumas, 2010; Gu et al., 2015; Sarmiento-Villamil et al., 2018).

An analysis of the corresponding OG0000022 protein-tree revealed that TFs harbouring the Ste12 DBD (IPR003120) are conserved even among the early-diverging fungal lineages such as the Chytridiomycota and Mucoromycota (**Supplementary item 10**). However, these were not detected among the plant-pathogenic fungi of the Ustilaginomycetes class in contrast to other Basidiomycetes such as *Puccinia striiformis*, where PstSTE12 is essential for pathogenicity on wheat (Zhu et al., 2018). In *Ustilago maydis*, a prominent maize pathogen of the Ustilaginomycetes, three other TFs were identified (Biz1, Mzr1 and Ztf1) that are reported to play related roles promoting infectious growth *in planta* (Flor-Parra et al., 2006; Zheng et al., 2008; Velez-Haro et al., 2020; de la Torre et al., 2020). These belong to a large clade of proteins restricted to the C2H2-DBD (**Supplementary item 10**) that included the BrlA conidiation regulators from the Aspergillaceae family and several regulators investigated in *M. oryzae* and *F. graminearum* with no obvious virulence-related function (Kwon et al., 2010; Son et al., 2011; M. Wang et al., 2015; Cao et al., 2016).

The common virulence-associated roles prompted a protein sequence alignment and phylogeny to visualise the relationship between characterised Ste12-orthologues and the *U. maydis* C2H2-domain regulators Biz1, Mzr1 and Ztf1 (**Fig. 6**). The highest-degree of conservation was observed among the Ste12-orthologues, and was centred at the corresponding DBD which did not extend to Biz1, Mzr1 and Ztf1. Although the C2H2-domains were largely conserved across both clades, other regions of significant identity were not obvious (**Fig. 6**). Hence, it is somewhat paradoxical that the sexual cycle is intimately connected with host invasion in *U. maydis* (Vollmeister et al., 2012), since both of these roles are driven by Ste12 in other fungi and the Ste12-DBD has been entirely lost in the Ustilaginomycete lineage. It is plausible that the C2H2-DBDs share a common evolutionary origin in this orthogroup, but is difficult to assess due to high-sequence divergence beyond this domain. Nevertheless, the evidence suggests a regulatory shift from Ste12 has occurred in *U. maydis*, possibly involving Biz1, Mzr1 and Ztf1, which share analogous roles promoting invasive growth. This is supported by findings that Biz1 and Ste12 are both activated and inhibited by related MAPK and cyclin-dependent kinase pathways respectively (de la Torre et al., 2020). The Ustilaginomycete class is almost entirely composed of plant-pathogens (Q.-M. Wang et al., 2015), indicating that Ste12-DBD loss could be a direct adaptation to host-driven selective pressures. Consequently, the functional relationships identified in this orthogroup analysis have presented an important hypothetical-link to the evolution of virulence.

**Fig. 6.**
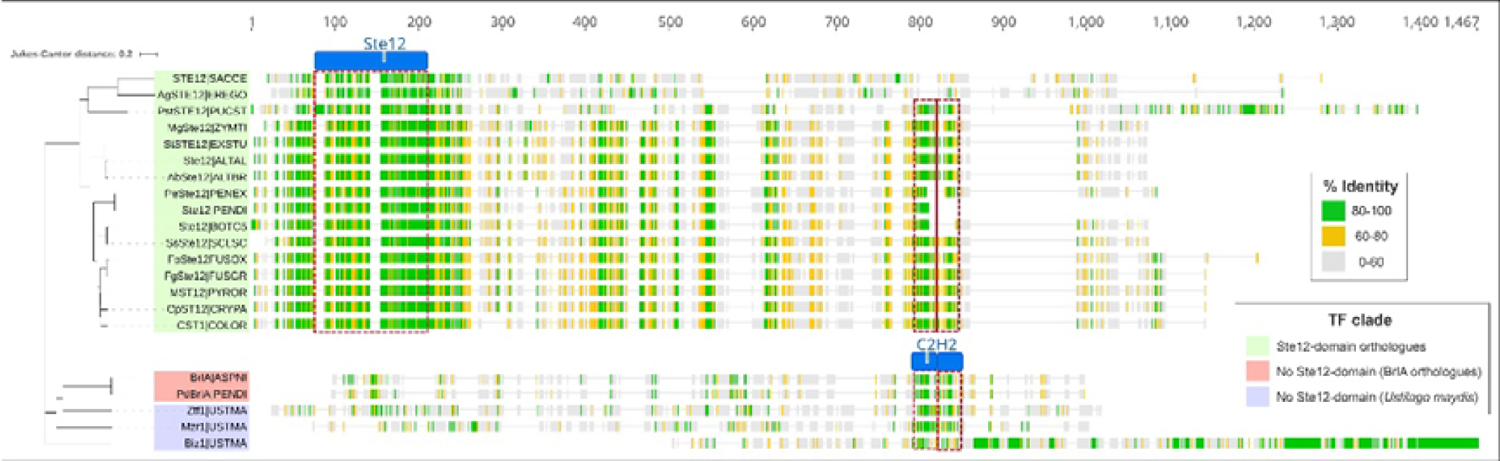
Characterised Ste12-orthologues relative to C2H2-only regulators of invasive growth A protein-sequence alignment is presented with a neighbour-joining tree depicting sequence divergence between Ste12-orthologues (highlighted in green), the *Ustilago maydis* C2H2-domain regulators of invasive growth (blue) and the Aspergillaceae BrlA regulators of conidiation (red) that stem from putative common lineages in orthogroup OG0000022. The protein names correspond to those reported in the relevant literature followed by the five-letter ORGIDs used in this study for the respective fungi. The alignment presents the % of residue identity across the aligned loci with gaps being ignored. The regions corresponding to the Ste12 (IPR003120) and the C2H2 (IPR013087) Interpro domains that were identified are presented above or within the alignments. Red boxes span the sequences where the respective domains are detected.

#### 3.3.3. Evidence for further expansion of the EBR1 TFs

The Zn2Cys6 TF EBR1 was first identified in *F. graminearum* through mutagenesis screening (Dufresne et al., 2008). The TF was subsequently investigated and shown to facilitate host invasion in *Fusarium* spp. by suppressing hyphal branching and promoting radial growth (Zhao et al., 2011). The large protein size and the internal positioning of the DBD, rather than the N-terminus, are relatively unusual features for the Zn2Cys6 TF family (MacPherson et al., 2006). In *F. oxysporum*, gene expansions are a characteristic of lineage-specific chromosomes (Coleman et al., 2009; Niño Sánchez et al., 2016). Interestingly, this phenomenon has been well-documented for *EBR1*, where as many as nine paralogues have been reported (Jonkers et al., 2014; van der Does et al., 2016). An analysis here of the corresponding OG0000121 protein-tree highlights cases in the different *F. oxysporum* isolates assessed here (**Supplementary item 11**), but also reveals EBR1 expansion across other important Ascomycete plant pathogens. This included the *M. oryzae* species-isolates assessed and the Mycosphaerellaceae fungal family. A protein sequence alignment/phylogenetic tree was therefore produced to investigate these putative expansions in greater detail (**Fig. 7**).

**Fig. 7.**
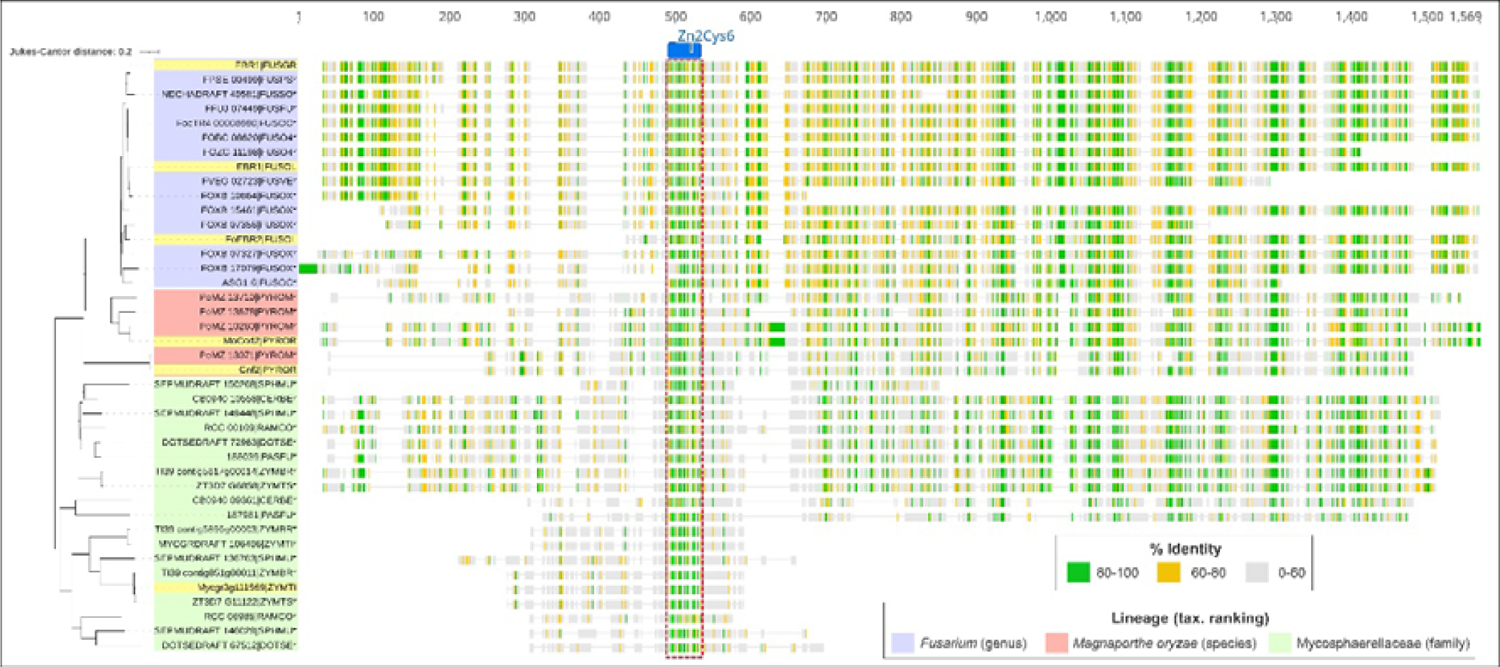
EBR1 expansion events identified in fungal pathogen lineages A protein-sequence alignment is presented with a neighbour-joining tree depicting sequence divergence between the EBR1-orthologues identified in orthogroup OG000121. The respective lineages (and their taxonomic ranking) are highlighted according to the figure legend, while yellow indicates those within each lineage that are characterised through functional investigation. The protein names correspond to those reported in the relevant literature followed by the five-letter ORGIDs used in this study for the respective fungi. The alignment presents the % of residue identity across the aligned loci with gaps being ignored. The region corresponding to the Zn2Cys6 (IPR001138) and Interpro domain that was identified is presented above the alignment. Red boxes span the sequences where the respective domains are detected.

In the *M. oryzae* reference isolate causing rice blast (PYROR) both MoCod2 (MGG_09263) and Cnf2 (MGG_15023) are TFs required for pathogenic development (Chung et al., 2013; Lu et al., 2014). The alignment revealed sequence conservation at the Zn2Cys6 DBD and several unannotated regions towards the C-terminus (**Fig. 7**). This strongly indicates both proteins stem from the EBR1 lineage, but from the fungi assessed here Cnf2 is unique to *M. oryzae* and has acquired an independent function in the virulence of this pathogen. The alignment also reveals that in the *M. oryzae* millet pathovar included in this study (PYROM), three MoCod2 paralogues exist (**Fig. 7**). The unique/redundant roles for these paralogues should be explored, as well as any functional interactions between MoCod2 and Cnf2 in *M. oryzae* to better understand the evolution and regulation of virulence by this TF lineage.

Several EBR1 duplication events were evident across the Mycosphaerellaceae family, which encompasses several damaging pathogens (**Supplementary item 11**). Inspection of the protein alignment (**Fig. 7**) suggests this is accompanied by the loss of a major proportion of the C-terminal conserved region in a sub-lineage which included *Z. tritici* Mycgr3g111569 (AlmA). This TF was characterised through overexpression which affected hyphal development during axenic growth (Cairns et al., 2015) and suggests a functional connection to other EBR1 orthologues is still maintained. Other TFs in the Mycosphaerellaceae-family lineage retain the conserved C-terminal region, but a functional role is yet to be reported.

It was striking that the majority of pathogens where EBR1 expansion events are observed are considered hemibiotrophs. It is plausible that a consequence of EBR1 duplication is the up-regulation of genes underpinning radial growth, thereby promoting asymptomatic colonisation of the host typical of this lifestyle. The novel evidence for gene expansions in key pathogen lineages, in particular the Mycosphaerellaceae, may warrant further exploration to establish their individual and/or connected roles during host infection.

#### 3.3.4. Tracing phytotoxic SM-gene cluster regulators

Pathway-specific regulators are typically encoded as part of fungal SM-gene clusters with most of those identified belonging to the Zn2Cys6 TF family (Keller, 2019; Lyu et al., 2020). A prominent feature of these clusters is diversifying selection and their frequent loss from fungal genomes, as well as horizontal transfer across taxa (Lind et al., 2017; de Jonge et al., 2018; Rokas et al., 2018). While a TF based analysis does not replace full SM-cluster phylogenies and functional investigation, a close inspection of the TF protein-tree clades revealed some surprising and significant connections for regulators of phytotoxic-SM biosynthesis.

An interesting case presented for BcBot6, a Zn2Cys6 TF assigned to orthogroup OG0000292 that regulates the biosynthesis of the phytotoxin botrydial in *Botrytis cinerea* (Porquier et al., 2016) (**Supplementary item 12**). A putative orthologue of BcBot6 was identified here in the phylogenetically-distant sunflower pathogen *Diaporthe helianthi* (DHEL01_v212770), and a sequence alignment with the adjacent branch on the protein-tree revealed little conservation relative to other fungal TFs (**Fig. 8A**). Although *D. helianthi* is reported to secrete several SMs that are phytotoxic (Ruocco et al., 2018), it is not known how these SMs are regulated, or if any are structurally analogous to botrydial. Moving forward, some interesting research questions that could be addressed are A) to explore the role of DHEL01_v212770 in *D. helianthi* or B) to cross-complement DHEL01_v212770 into the *bcbot6* deletion mutants of *B. cinerea* to determine if it is functionally conserved. The sequence alignment also suggested the putative BcBot6 orthologue annotated in the T4 isolate of *B. cinerea* (BOTC4) was truncated relative to BcBot6 in *B. cinerea* B05.10 (BOTC5) (**Fig. 8A**). Nevertheless, botrydial is an important virulence factor produced by T4 (Pinedo et al., 2008; Porquier et al., 2016) and further analysis will be required to define translational or regulatory differences attributable to this difference.

**Fig. 8.**
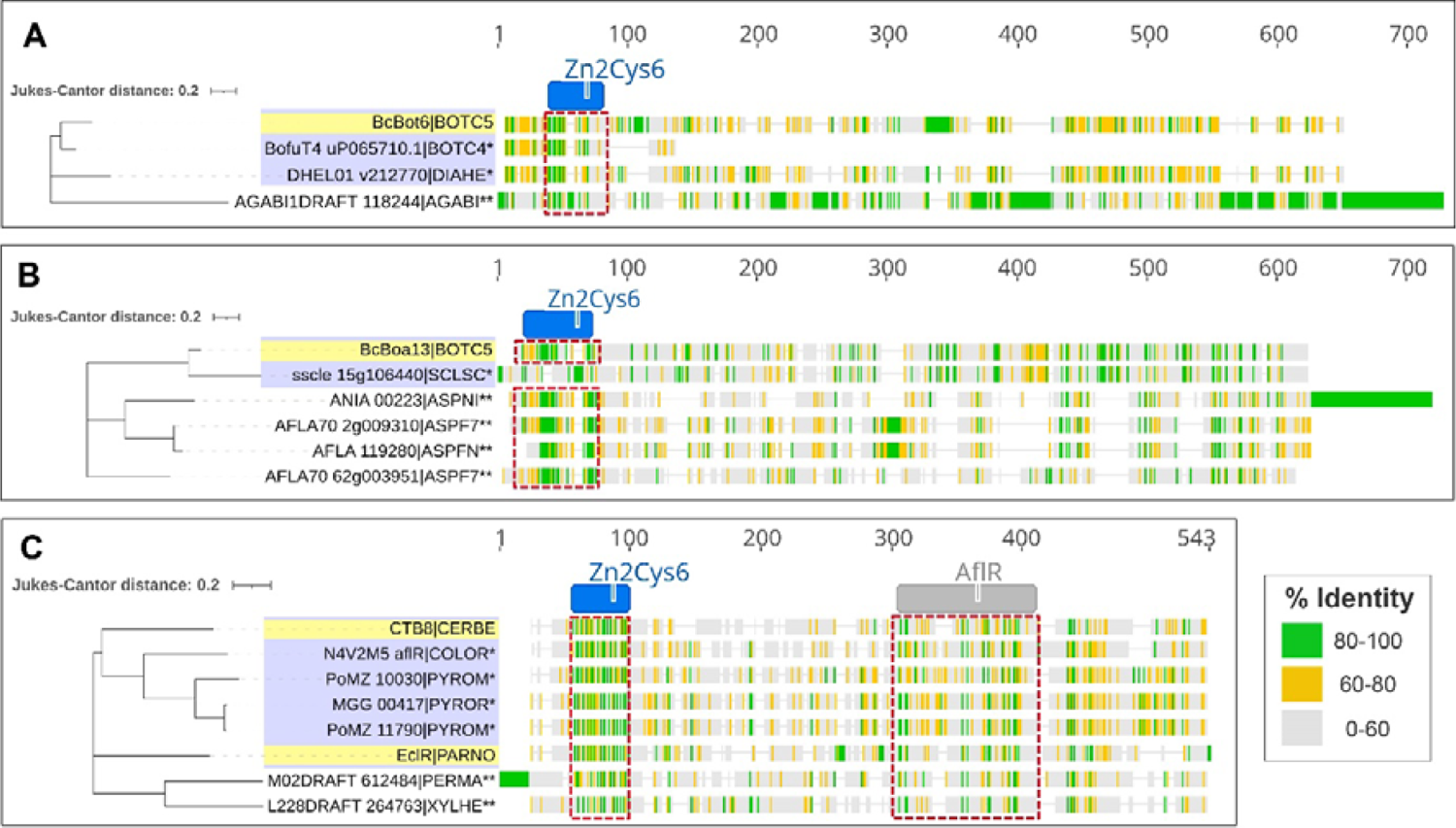
Phytotoxic secondary-metabolite cluster regulators across fungal pathogen lineages Protein-sequence alignments are presented with a neighbour-joining tree depicting sequence divergence between A) the *B. cinerea* BcBot6 regulator of botrydial biosynthesis, B) the BcBoa13 regulator of botcinic acid biosynthesis and C) the CTB8 regulator of cercosporin biosynthesis. Putative homologous-regulators are highlighted in blue, while yellow indicates those within each lineage that are characterised through functional investigation. The protein names correspond to those reported in the relevant literature followed by the five-letter ORGIDs used in this study for the respective fungi. The alignment presents the % of residue identity across the aligned loci with gaps being ignored. The region corresponding to the Zn2Cys6 (IPR001138) and (in panel C only) the AflR (IPR013700) Interpro domains that were identified are presented above the alignment. Red boxes span the sequences where the respective domains are detected. Homologous proteins that are yet to be functionally verified are indicated with by the asterix* and the double asterix** indicates adjacent clades in the respective protein-trees (**Supplementary items 12 & 13**) that are not predicted to encode functional homologues.

Another *B. cinerea* pathway-specific phytotoxin regulator is BcBOA13, which controls botcinic acid biosynthesis and is present in the B05.10 isolate but missing in the T4 isolate (Dalmais et al., 2011; Porquier et al., 2019). Interestingly, a homologous botcinic acid biosynthetic gene-cluster was previously identified in the genome of *Sclerotinia sclerotiorum*, a broad-host range pathogen that is closely related to *B. cinerea* (Porquier et al., 2019). However, inspection of the BcBOA13 clade in the OG0000000 orthogroup protein-tree revealed the homologous protein sscle_15g106440 was absent (**Supplementary item 12**). A closer inspection and sequence alignment that included sscle_15g106440 (Uniprot accession A0A1D9QLR3) suggested a replacement has occurred spanning the N-terminal DBD (**Fig. 8B**). It will be important to ascertain whether the sscle_15g106440 protein-annotation is accurate, or whether expression of the SM-gene cluster in *S. sclerotiorum* has been affected due to this mutation. The nearest branches in the BcBOA13 clade correspond to TF homologues that regulate the biosynthesis of aspirochlorine, a mycotoxin produced in *Aspergillus* spp. with no structural or functional relationship to botcinic acid having been described (Chankhamjon et al., 2014; Uka et al., 2020). Hence, it is unlikely that a BcBOA13 biosynthetic cluster was horizontally transferred across these fungal genomes. If a common ancestral protein existed, the sequence alignment indicates significant divergence has occurred relative to BcBOA13 outside the Zn2Cys6-DBD and some C-terminal residues (**Fig. 8B**).

Cercosporin and its structural homologues are potent light-activated phytotoxic SMs produced by *Cercospora* spp. (Daub et al., 2005; Newman and Townsend, 2016). Evidence of horizontal transfer, lineage diversification and/or gene loss has recently been detailed for the cercosporin biosynthetic-gene cluster in several Ascomycete lineages (de Jonge et al., 2018). *CbCTB8* encodes a cercosporin pathway-specific Zn2Cys6 regulator in *Cercospora beticola* and several cluster-orthologues were identified here in the protein-tree clade (**Supplementary item 13**). This included *Parastagonospora nodorum* EclR, a TF that controls the production of the phytotoxic structural homologue of cercosporin, elsinichrome C (Chooi et al., 2017). A sequence alignment was therefore undertaken to explore the conserved protein regions in CbCTB8, EclR and the other predicted cercosporin-regulators that were identified in *M. oryzae* pathovars and the cucurbit pathogen *Colletotrichum orbiculare* (**Fig. 8C**). The alignment revealed that, in addition to the N-terminal Zn2Cys6 DBD, the AflR domain associated with regulators of aflatoxin biosynthesis (Yu et al., 1996; Liu and Chu, 1998) is co-conserved. This was also observed in the adjacent protein-tree clade that included TFs from the symbiotic endophytes *Xylona heveae* and *Periconia macrospinosa*, suggesting the domains are not strictly-linked to the control of phytotoxin production (**Fig. 8C**). The distinguishing features of CbCTB8 orthologues that target genes controlling phytotoxin biosynthesis must therefore be determined, yet their common evolutionary lineage was predicted in line with the more comprehensive gene-cluster analysis conducted previously (de Jonge et al., 2018).

## 4. Conclusions

The aims of the analysis presented here were to explore the conservation, expansion and loss of TFs associated with pathogenic lifestyles, as well as provide a novel resource to effectively trace the evolutionary trajectory of regulators that are correlated with virulence. A taxonomically-diverse range of fungi were assessed and a significantly-reduced TF content was observed in obligate host-associated pathogens. This suggested that, in general, the limited ecological-niche of these fungi lends itself to regulation by relatively few TFs. This study then resolved the TF classification system from 70 DBD-based families to an informative set of 855 orthogroups. Recent approaches have used large genomic datasets to demonstrate that the content of other distinct gene families, such as carbohydrate-acting enzymes, can both predict and refine our definitions of plant-pathogenic lifestyles (Hane et al., 2020; Haridas et al., 2020). These definitions more accurately reflect the molecular factors that underpin virulence. Here we demonstrated that many TF orthogroups were over/underrepresented in distinct pathogen lineages (**Fig. 4**). This suggests accurate information on both the type and content of fungal TFs should complement these approaches and enhance our understanding of the factors that contribute to virulence in plant-pathogenic fungi.

For TFs that have been under strong diversifying selection, precisely inferring their true orthologues will require detailed datasets that can incorporate a large number of closely related species. Nevertheless, as demonstrated in the virulence regulators discussed, the exploration of specific clades within the orthogroup protein-trees has provided novel insights into their evolutionary origins and trajectories. Importantly, the orthogroups presented in this analysis are a useful resource for the future exploration of virulence-regulating TFs. It is envisaged this will occur using different and complementary approaches:

- The orthogroup lineages (protein-trees) of functionally characterised TFs can be explored to define taxonomic orthologues and search for cases of gene duplication or loss in a pathogen of interest.
- Investigate orthogroups that exhibit expansion in a pathogen lineage. These can be identified from the heatmap in **Fig. 4 & Supplementary item 7** which correspond to columnar layout of the orthogroup counts in **Supplementary item 6**. The protein-trees for the expanded orthogroups can then be explored in detail to guide further phylogenetic and functional analyses that investigate any lifestyle-associated selection pressures.
- Search for cases of horizontal-gene transfer events for TFs implicated in the acquisition of fungal virulence. The orthogroup clade of a known virulence regulator can be examined in the corresponding protein-tree to identify anomalous TF neighbours (i.e. those from taxonomically distant fungi). This is particularly relevant for regulators of phytotoxic SM-gene clusters where genetic mobility is common (Lind et al., 2017; de Jonge et al., 2018; Rokas et al., 2018).

Combined, these are powerful means to assist the characterisation of the fundamental virulence regulators in plant-pathogenic fungi. This will provide insight into the distinct factors underpinning plant infection to assist the design of targeted mitigation strategies. This could include the direct inhibition of the TFs themselves, or their upstream/downstream signalling components.

## Supporting information

Supplemental item 1

Supplemental item 2

Supplemental item 3

Supplemental item 4

Supplemental item 5

Supplemental item 6

Supplemental item 7

Supplemental item 9

Supplemental item 10

Supplemental item 11

Supplemental item 12

Supplemental item 13

## Acknowledgements

We would like to that Dr. Mark Derbyshire, Dr. Robert Syme, Dr. Darcy Jones and Dr. James Hane for their advice and support in undertaking the computational analysis.

## Funding

This study was supported by the Centre for Crop and Disease Management, a joint initiative of Curtin University (https://www.curtin.edu.au/) and the Grains Research and Development Corporation (https://grdc.com.au/) [research grant CUR00023]. Evan John was supported by an Australian Government Research Training Program Scholarship (https://www.dese.gov.au/) administered through Curtin University. This work was also supported by resources provided by the Pawsey Supercomputing Centre with funding from the Australian Government and the Government of Western Australia. The funders had no role in study design, data collection and analysis, decision to publish, or preparation of the manuscript.

## Supplementary items

**Supplementary item 1 - Organism and TF annotation metadata**

A table providing the fungal species/strains used in the analysis [columns 1-2], their assigned five letter ORGIDs [3], alternative names used for each organism in the scientific literature [4], their lifestyle (either necrotroph, biotroph, hemibiotroph, symbiont or saprophyte) [5], host association (either facultative or obligate and not applicable for saprophytes) [6], TF count relative to the proteome size [7-9], details on the proteome sources [10-11], references for the associated genome publication [12-13] and the NCBI taxonomic identifiers with phylogenetic rankings for each organism [14-20].

**Supplementary item 2 - Assessment of total TF DBDs compiled**

A table listing the respective Interpro IDs [column 1], Pfam/Superfam IDs [2], DBD/TF family description [3], the total count across species analysed in this study [4] and an indication whether the DBD was in the list adapted for this study [5] (Shelest, 2017).

**Supplementary item 3 - Orthogroups defined for fungal TFs**

Spreadsheets detailing individual TFs assigned to orthogroups (spreadsheet 1) or the unassigned TFs (spreadsheet 2). Orthogroups are listed by size in descending order. TF identifiers follow the format: ‘ProteinID | ORGID’ where the ProteinID was derived from the downloaded proteome source.

**Supplementary item 4 - TF-distance based species-tree**

The species-tree built from orthogroup TF distances (Emms and Kelly, 2018) and rooted at *Synchytrium endobioticum* (SYNEN) which was used for orthologue inferences. The full fungal species/strain IDs are included alongside the respective ORGIDs at the tree leaves.

**Supplementary item 5 - NCBI taxonomy tree**

The NCBI taxonomy tree for the fungal species/strains used in the analysis. Node points indicate the respective taxonomic divisions listed in **Supplementary item 6**. The full fungal species/strain IDs are included alongside the respective ORGIDs at the tree leaves.

**Supplementary item 6 - TF orthogroup counts**

A spreadsheet detailing the orthogroup counts for each organism [column 1], which are listed using their ORGIDs in the order of the leaf nodes in the TF species-tree. The respective NCBI taxonomic rankings for species, genus, family, order, class and phylum are listed [2-7]. The listed TFs for 855 orthogroups [8-862] are ordered by the hierarchical clustering depicted in **Fig. 4**.

**Supplementary item 7 - Clustered TF-orthogroup heatmeap (high resolution)**

A high-resolution heatmap of normalised counts (Z-scores) for the 855 hierarchically-clustered orthogroups as depicted in **Fig. 4**. Fungal ORGIDs are listed on the Y-axis and TF-orthogroup IDs are listed on the X-axis which correspond to the layout for the TF counts presented in **Supplementary item 6**.

**Supplementary item 8 - TF orthogroup protein-trees**

Accessible via: https://doi.org//10.6084/m9.figshare.17211737

The TF orthogroup protein-trees generated in this study (for the orthogroups OG0000000 > OG0000591). These are in Newick format and can be visualised in the reader’s chosen tree-viewing platform such as iTOL (Letunic and Bork, 2019). The branch lengths are defined by the DendroBLAST distance measure (Kelly and Maini, 2013).

**Supplementary item 9 - OG0000017 protein-tree**

The full OG0000017 (Pf2-orthologue) protein-tree harbouring the TFs explored in **Fig. 5**. Highlighted are the *S. cerevisiae* MAL13 outgroup in blue, *A. nidulans* AmyR in orange and the Pf2 taxonomic clade (green) with the functionally defined orthologues in yellow (for the fungi FUSVE, FUSGR, NEUCR, PYROR, BOTC5, ALTBR, PYRTR, PARNO and ZYMTI respectively). The branch length is defined by the DendroBLAST distance measure (Kelly and Maini, 2013).

**Supplementary item 10 - OG0000022 protein-tree**

The full OG0000022 (Ste12) protein-tree harbouring the TFs explored in **Fig. 6**. Highlighted are the Ste12 orthologues in green with the functionally defined TFs in yellow (for the fungi ZYMTI, ALTBR, ALTAL, EXSTU, PENDI, PENEX, SCLSC, BOTC5, FUSGR, FUSOX, PYROR, CRYPA, VERDA, COLOR and PUCST respectively) and the Ste12 TFs that lack C-terminal C2H2 domains in blue (for SACCE and EREGO). The *U. maydis* C2H2-only TFs Biz1, Mrz1 and Ztf1 are highlighted in purple and the BrlA orthologues (PENDI and ASPNI) in orange. The branch length is defined by the DendroBLAST distance measure (Kelly and Maini, 2013).

**Supplementary item 11 - OG0000121 protein-tree**

The full OG0000121 (EBR1) protein-tree harbouring the TFs explored in **Fig. 7**. Highlighted are the functionally defined EBR1 orthologues in yellow (for the fungi FUSOL, FUSGR, ZYMTI and PYROR respectively) and the putative expansions in the *Fusarium*, *M. oryzae* and the Mycosphaerellaceae lineages. The branch length is defined by the DendroBLAST distance measure (Kelly and Maini, 2013).

**Supplementary item 12 - OG0000292 (BcBot6) and OG0000000 (node 6240 - BcBOA13) protein-trees**

Panel A highlights the BcBot6 transcription factor (TF) for *B. cinerea* T4 (BOTC4) and B05.10 (BOTC5) isolates in yellow alongside the putative *D. helianthi* (DIAHE) homologue in blue, indicating a shared ancestral origin despite relatively large species distances. Panel B highlights the BcBOA13 TF from BOTC5 in yellow and the aspirochlorine TF regulators from *Aspergillus* spp. (corresponding to ORGIDs in blue: ASPNI, ASPFN and ASPF7) that also suggest a shared ancestral origin despite distant fungal taxonomies. The branch length is defined by the DendroBLAST distance measure (Kelly and Maini, 2013).

**Supplementary item 13 - OG0000000 protein-tree (node 6486 - CbCTB8)**

A tree representing the relationship between transcription factors regulating the biosynthesis of Cercosporin (i.e. CbCTB8) or its structural homologues. The functionally defined CbCTB8 orthologues in the literature (represented by the fungal ORGIDs for CERBE and PARNO) are highlighted in yellow. Other putative homologous regulators reported for the cercosporin secondary-metabolite-gene clusters (de Jonge et al., 2018) were also grouped in a neighbouring orthogroup-clade and are highlighted in blue (represented by the fungal ORGIDs for PYROM, PYROR and COLOR). The branch length is defined by the DendroBLAST distance measure (Kelly and Maini, 2013).

## Abbreviations

TF: Transcription factor

DBD: DNA-binding domain

TFome: complete TF repertoire

SM: secondary metabolite

## Notes

https://doi.org//10.6084/m9.figshare.17211737

### Competing Interest Statement

The authors have declared no competing interest.

