## Supplementary figures and images for "Transcription factor lineages in plant-pathogenic fungi, connecting diversity with fungal virulence"

### Supplemental item 4

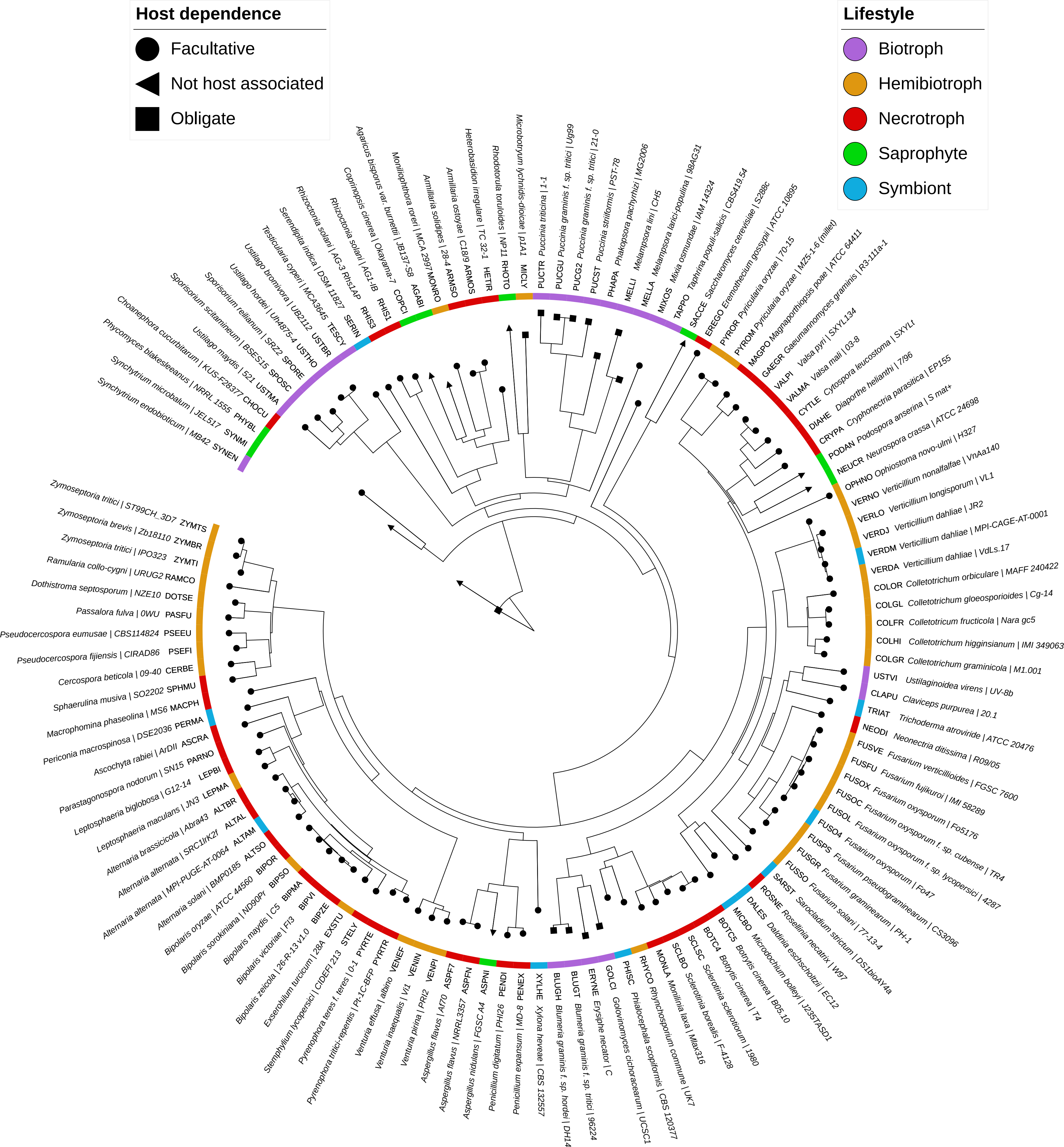

### Supplemental item 5

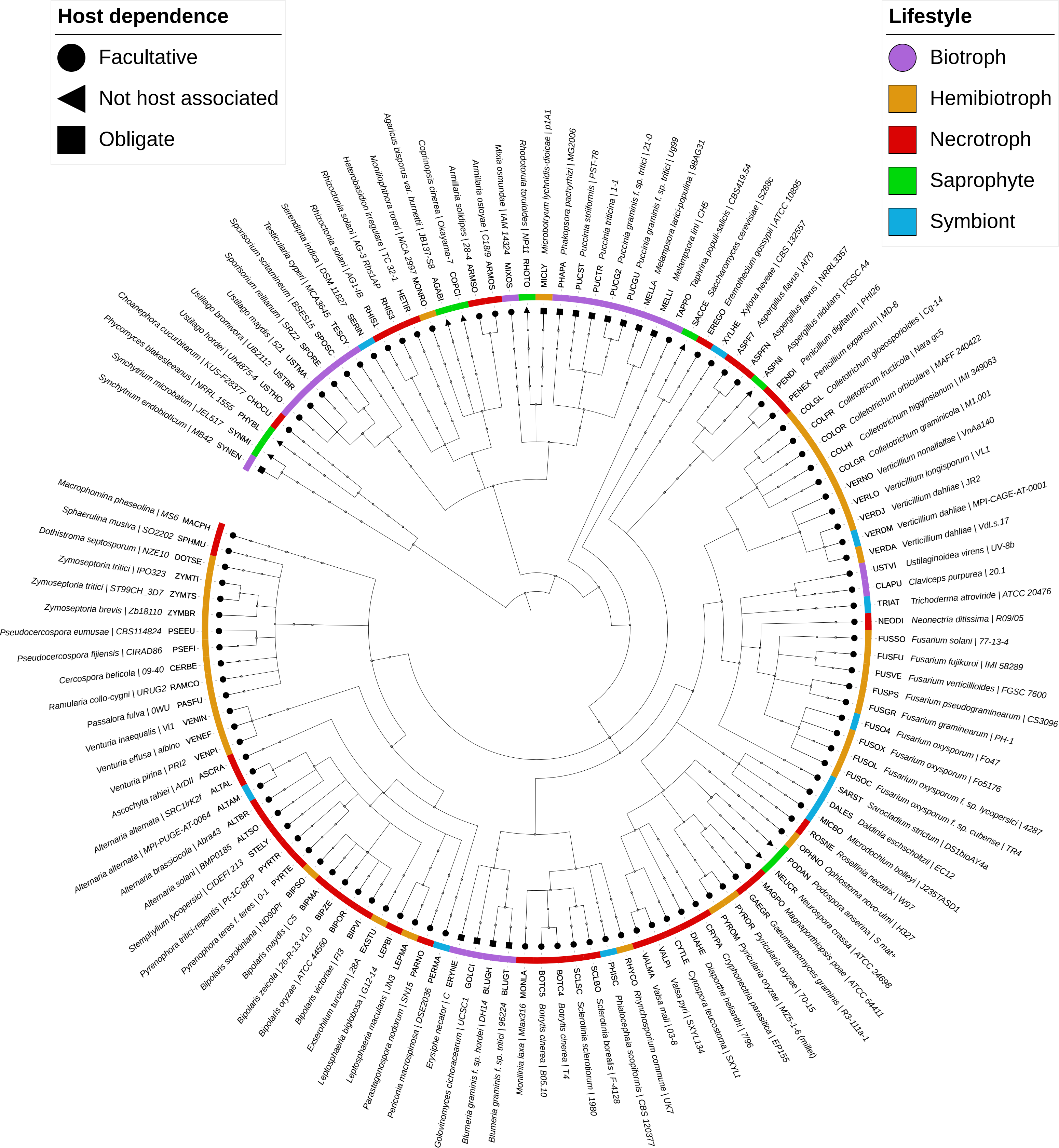

### Supplemental item 7

Clustered fungal transcription factor orthogroups

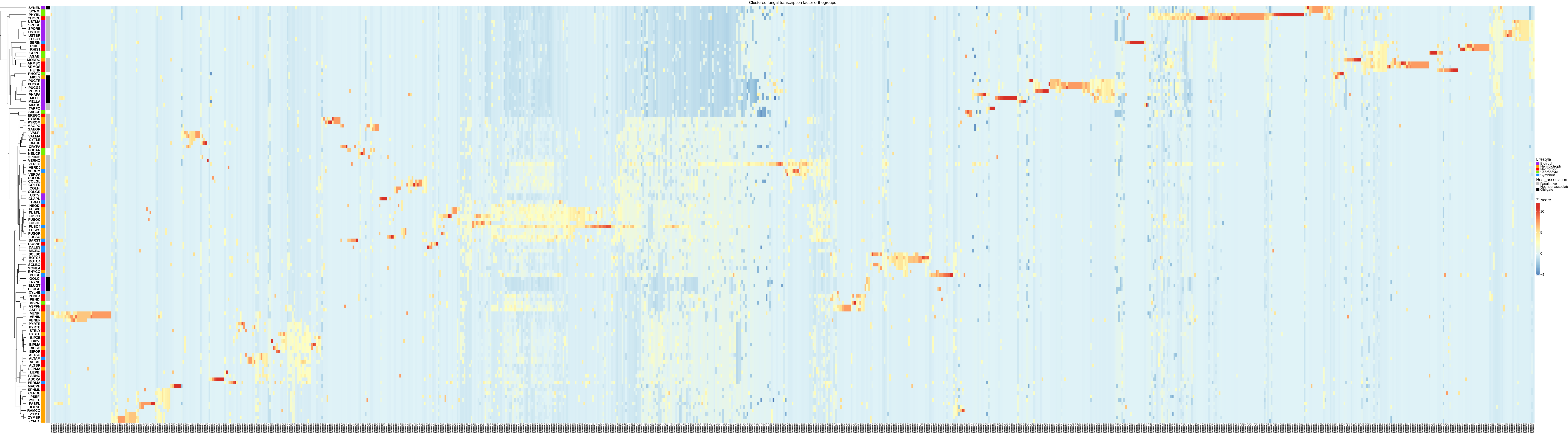

### Supplemental item 9

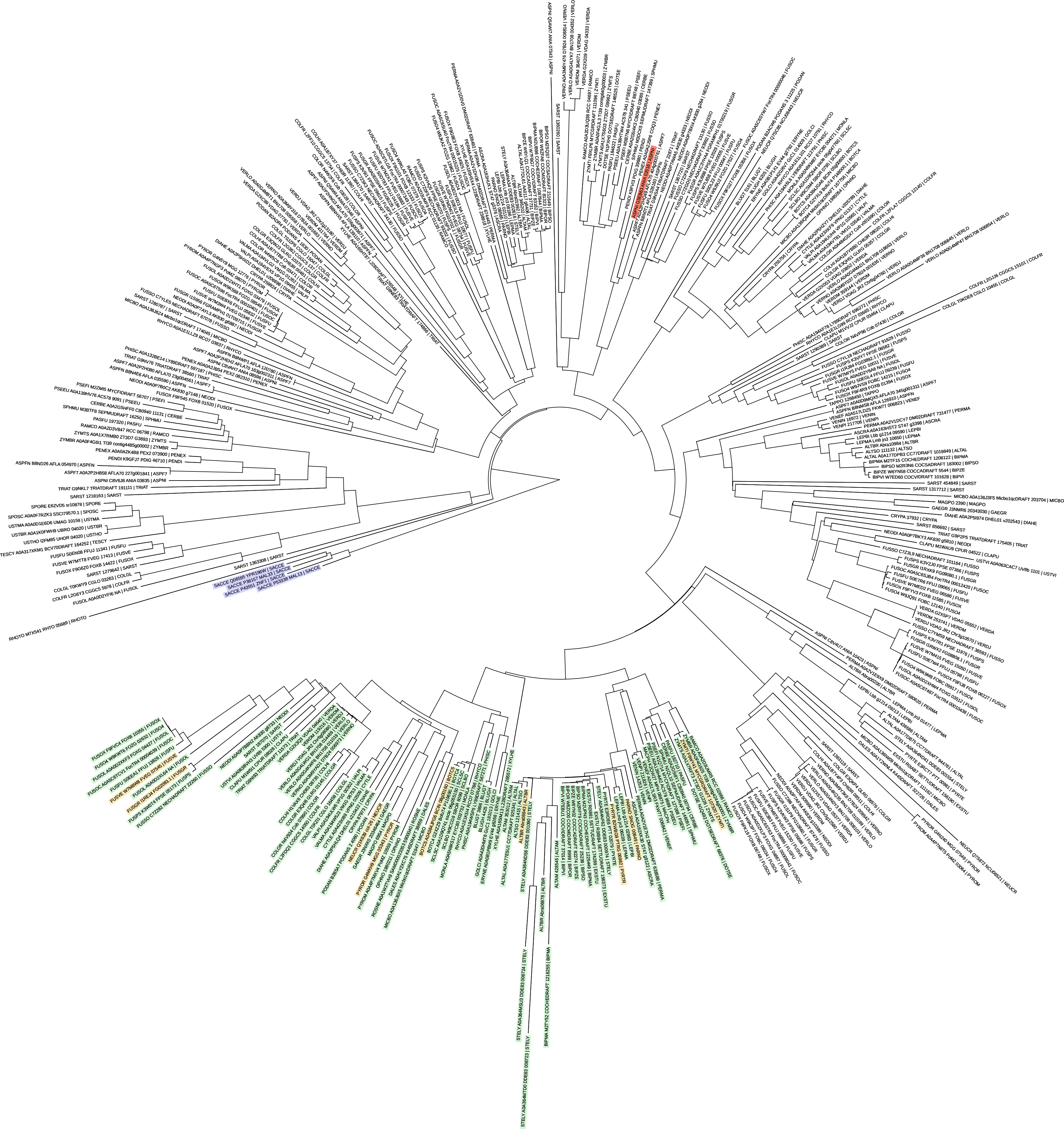

### Supplemental item 10

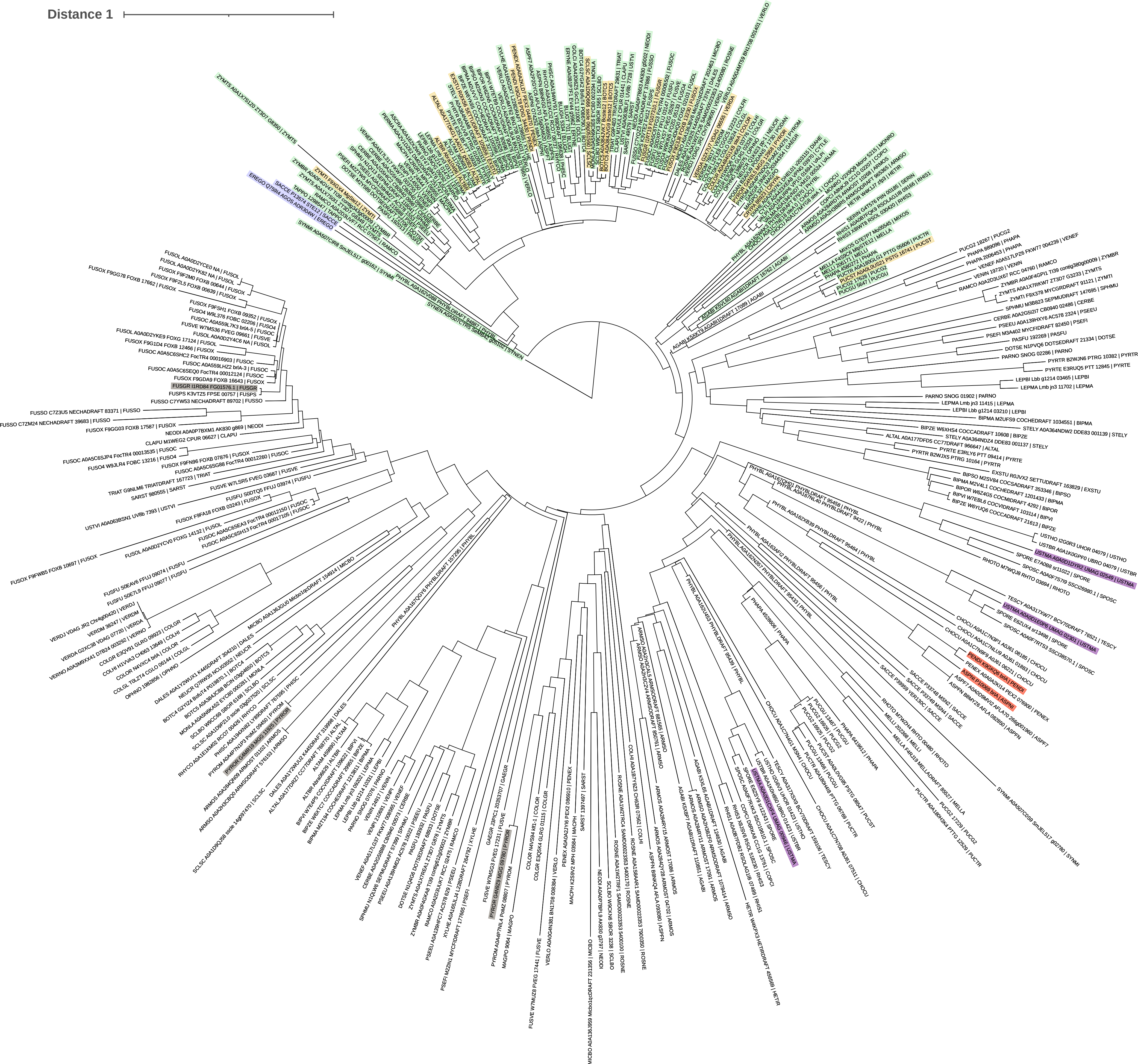

### Supplemental item 11

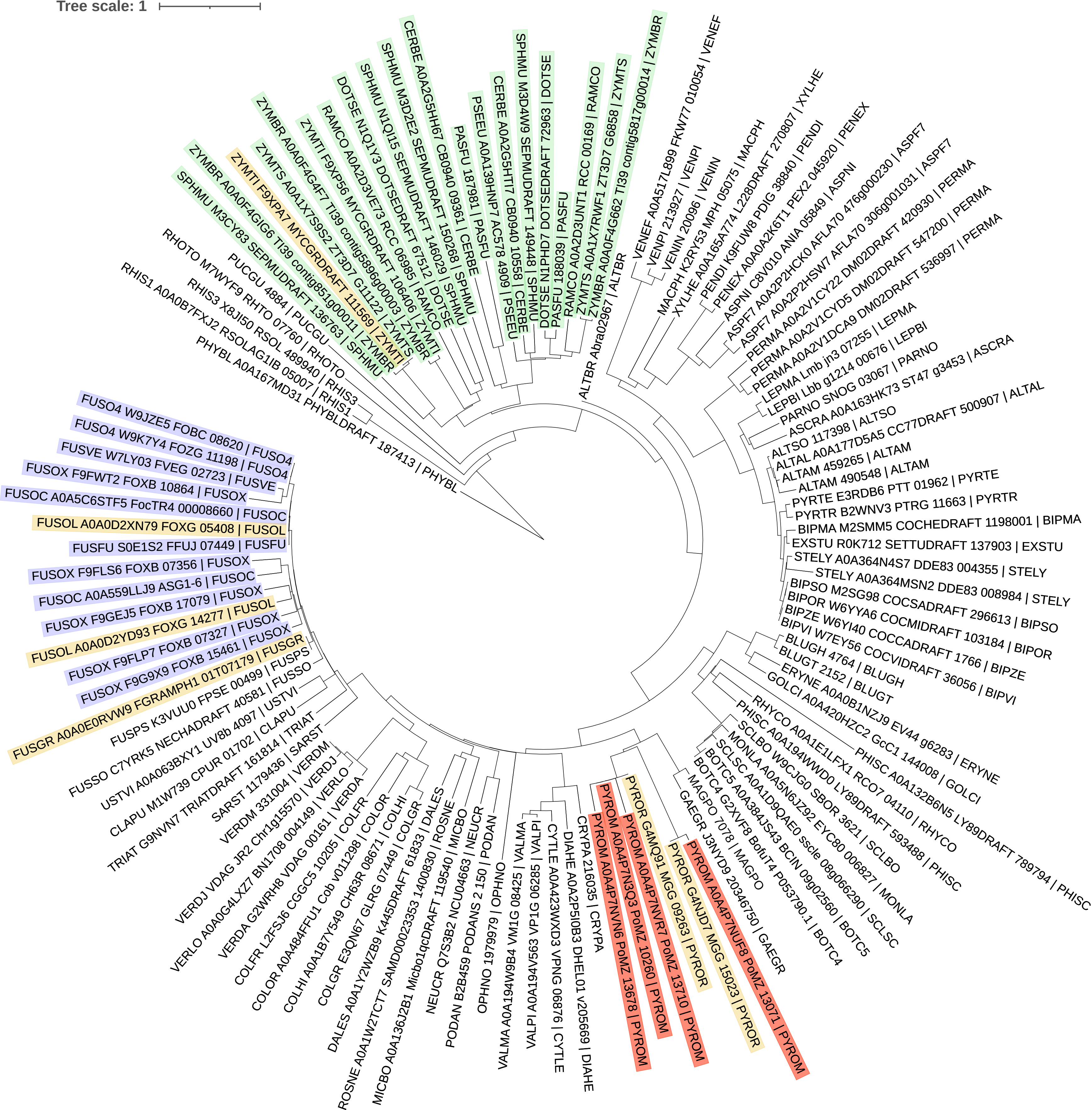

### Supplemental item 12

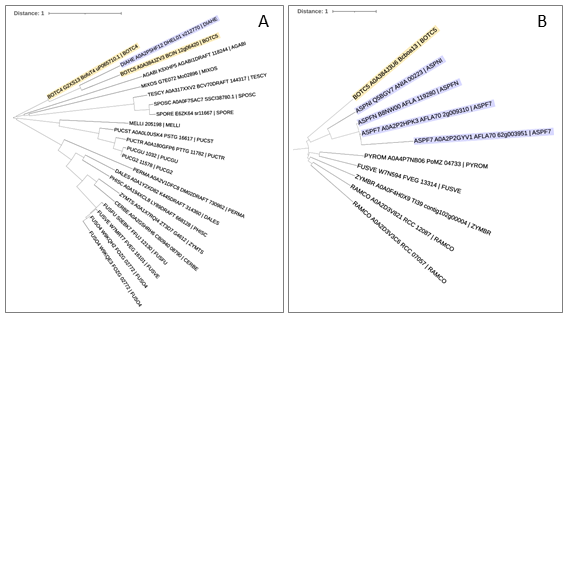

### Supplemental item 13

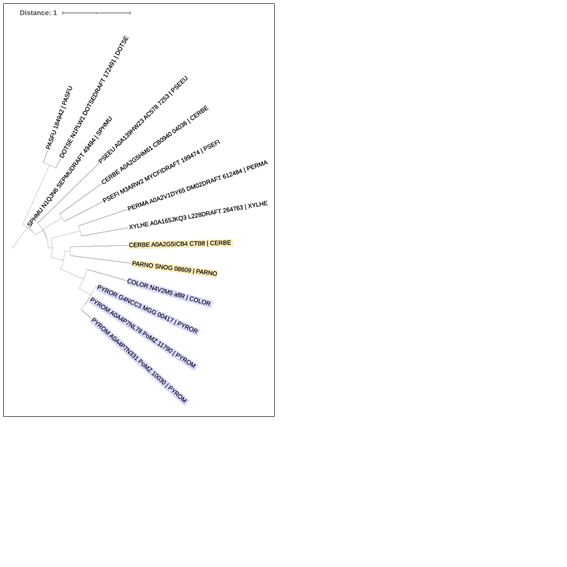
